# A multi-model Framework for the Arabidopsis life cycle

**DOI:** 10.1101/358408

**Authors:** Argyris Zardilis, Alastair Hume, Andrew J. Millar

## Abstract

Linking our understanding of biological processes at different scales is a major conceptual challenge in biology, which is aggravated by differences in research methods. Modelling can be a useful approach to consolidating our understanding across traditional research domains. The laboratory model species *Arabidopsis thaliana* is very widely used to study plant growth processes and has also been tested more recently in eco-physiology and population genetics. However, approaches from crop modelling that might link these domains are rarely applied to Arabidopsis. Here, we combine plant growth models with phenology models from eco-physiology, using the agent-based modelling language Chromar. We introduce a simpler Framework Model of vegetative growth for Arabidopsis, *FM-lite.* By extending this model to include inflorescence and fruit growth and seed dormancy, we present a whole-life-cycle, multi-model *FM-life,* which allows us to simulate at the population level in various genotype x environment scenarios. Environmental effects on plant growth distinguish between the simulated life history strategies that were compatible with previously-described Arabidopsis phenology. Our results simulate reproductive success that is founded on the broad range of physiological processes familiar from crop models and suggest an approach to simulate evolution directly in future.

**Highlight:** A whole-life-cycle multi-model for *Arabidopsis thaliana* combines phenology and physical growth models to explain reproductive success in different genotype x environment scenarios.

## Introduction

Understanding the links between biological processes at multiple scales, from molecular regulation to populations and evolution, is a major challenge in understanding life. As systems become more complex we need models to describe our understanding and help our thinking when trying to explain and make predictions across scales. This approach has also been proposed in attempts to engineer crop traits starting from genetics or from genomes (Welch *et al.*, 2005; Yin and Struik, 2008, 2010; Parent and Tardieu, 2014; Wu *et al.*, 2016; Chenu *et al.*, 2018), where simpler models have demonstrated both the potential of crop modelling in general and the significant demands of detailed models for empirical data that varies in availability (Hammer *et al.*, 2006; Asseng *et al.*, 2013). For microorganisms, comprehensive models link the metabolic and molecular level with the cellular (Karr *et al.*, 2012) and population growth scales (Weiße *et al.*, 2015), whereas contemporary work in more complex organisms has necessarily focused more narrowly (Buckley and Mott, 2013; Lynch, 2013; Zhu *et al.*, 2013; Klose *et al.*, 2015; Le Novere, 2015; Hepworth et *al.,* 2018).

The concentration of plant science research on the laboratory model species *Arabidopsis thaliana* offers an opportunity for broad understanding that includes mechanistic models (Chew et al., 2014a; Voss et al., 2014, Urquiza et al., this volume). The Framework Model (FMv1) represented vegetative growth of Arabidopsis in lab conditions (Chew *et al.*, 2014b), starting from four independent models that represent photosynthesis and carbon storage (Rasse and Tocquin, 2006), plant structure and carbon partitioning among organs (Christophe *et al.*, 2008), flowering phenology (Chew *et al.*, 2012) and the circadian clock gene circuit and its output to photoperiodic flowering (Salazar *et al.*, 2009). Later updates focussed on plant phenotypes controlled by the clock, such as tissue elongation and starch metabolism (FMv2; Chew *et al.*, 2017), or temperature and organ-specific inputs to flowering (Kinmonth-Schultz *et al.*, 2018). The Framework Models align with community efforts to link understanding of crop plant processes at multiple scales, for benefits in agriculture (Wu *et al.*, 2016; Zhu *et al.*, 2016). Among the limitations of the Framework Models, growth was limited to the vegetative stage, ending upon flower induction. Without reproduction, the models had no seed yield or link to evolutionary fitness. Without seed dormancy, they lacked a major determinant of Arabidopsis life history in the field. Their representation of the circadian clock was also unnecessarily detailed for many studies outside chronobiology.

Other models have considered reproductive success through growth, including for Arabidopsis. One simplified approach relates growth and fitness only to the duration of the developmental period and not to its timing in the year, ignoring environmental influences (Prusinkiewicz *et al.*, 2007). On the other hand, ecological phenology models of the Arabidopsis life cycle consider natural environmental conditions but ignore physical growth and development (Chuine and Beaubien, 2001; Chuine, 2010; Burghardt *et al.*, 2015). We apply a declarative agent-based modelling approach (Urquiza et al., this volume) that facilitates model composition, to develop FM-life, an extension of the Framework Model to the whole Arabidopsis life cycle. FM-life includes a simpler model of vegetative growth, FM-lite, without the clock circuit, and a new model of inflorescence growth including reproduction. We introduce a clustering approximation, in order to simulate FM-life tractably at the population scale over decades. Testing the FM-life model with contrasting environmental and genetic inputs shows that ecological questions can increasingly be informed by mechanistic understanding of growth processes (Millar, 2016; Doebeli *et al.*, 2017).

## Materials and Methods

### Chromar

FMv1 and FMv2 are both available as Matlab programs. Here we use a declarative, agent-based language Chromar, which represents the models very concisely and supports simulations (Honorato-Zimmer *et al.*, 2017). Chromar models are designed to be human-readable. As no agent-based approach is broadly familiar in biological modelling, we introduce simple examples below. Briefly, agents in Chromar conform to a small number of types. Each type defines the attributes of the agent. A type defined as Leaf(mass: real) represents leaves with a mass attribute that is a real number, for example Leaf(mass = 5). Agents operate stochastically, according to rules that specify:

- Agent level dynamics, where agents are created or destroyed. For example, organogenesis of a leaf with mass *m_0_* at rate *k* corresponds to the following rule: 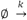 Leaf (mass = *m*_0_)
- Attribute level dynamics, which alter the attributes of an agent. For example, growing a leaf by mass *g* at rate *k_g_* corresponds to: Leaf(mass = *m*) 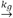 Leaf(mass = *m + g*)

Two further features of Chromar are particularly useful *here, fluents* for describing time-dependent, deterministically changing values such as environmental inputs, and *observables* used for capturing system-wide state easily. Observables are used to manage multiple descriptions of the same process at different levels of abstraction, such as the individual leaves of a plant and the total number of leaves in a rosette. Fluents and observables can be used directly in expressions and be combined with the normal set of mathematical expressions, so we could have a function, *f(n,* temp), that takes as arguments and applies some mathematical operation to an observable for the number of leaves, *n,* and a fluent, temp, for the temperature.

Here we made use of a particular implementation of Chromar as an embedded domain specific language inside the general purpose programming language Haskell (Honorato-Zimmer *et al.*, 2018). This means that we can leverage the power of a general-purpose programming language inside the rules, for example for describing attribute dynamics or rule rates.

### Phenology models in Chromar

In many phenology models, the simulated plant accumulates a conceptual development indicator in every time unit as a function of the contributing environmental factors, until a threshold is reached for transition to the next developmental stage. For example, in a seed type Seed(dev: real), the dev attribute measures development towards germination. A phenology rule for germination affected by temperature and moisture, starting from dev value *d,* could be:

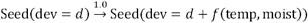

On average once every time unit the dev attribute of a particular seed will be increased from the present value, *d,* by a function of the contributing factors temp and moist. Further parameters might represent how sensitive the seed is to the environmental factors. At the threshold *D_t_,* the seed germinates to a plant and resets the development measure to 0:

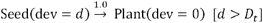

where the expression inside the square brackets is used to indicate conditional activity of the rule. The rule is active only when the expression evaluates to true.

### The component models

The models presented here represent the full life cycle in three stages: seed dormancy (A, left panel, Figure 1), vegetative growth up to flowering (B, left panel, Figure 1), and the reproductive stage up to seed dispersal (C, left panel, Figure 1). Each model (A, B, C) includes a phenology component that represents only timing (Section 2.2). The vegetative and reproductive stage models also represent biomass growth at the organ level, based on the carbon budget of the plant. We varied genetic parameters that affect only the timing components of A (seed dormancy, *ψ_i_*) and B (floral repression during vegetative growth, *fi),* for comparison to (Burghardt *et al* 2015). Each parameter value for an individual plant can be fixed or selected probabilistically from a distribution as described (Burghardt *et al.*, 2015). The three models were integrated in a whole life-cycle model of one plant (FM-life), and then extended to a population of such plants.

**Figure 1.**
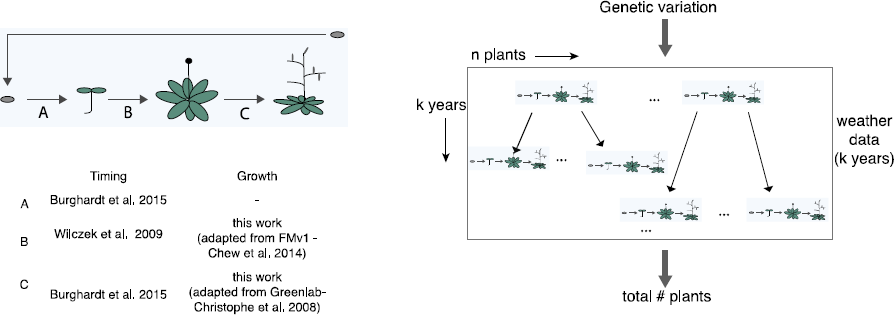
Overview of the FM-life and population models used in this study. Left Overview of the models used for the growth and timing components for the three developmental stages: seed dormancy (A), vegetative period (B), and reproductive period (C) and Right Sketch of the population level model. Inputs to the model are the distribution of values of the two genetic parameters (f, ψ and weather data from some location for a number of years. The output is some population measure of interest, an example might be the total number of plants after k years.

### Seed dormancy model (A)

The seed dormancy model is the Chromar version of the model of (Burghardt *et al.*, 2015), which is based in turn on (Alvarado and Bradford, 2002). It represents the development of a newly-dispersed seed from dev = 0 to a threshold value, *D_g_*, where the seed germinates. Above baseline levels of temperature *T_b_* and of moisture (see below), increasing moisture and temperature speed the progress towards germination. The additional developmental units added (hydrothermal units, htu) at every time unit are described by:

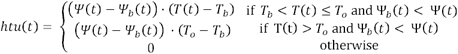

where *Ψ*(*t*) and *T(t)* give the moisture and temperature levels at time *t* respectively. The definition distinguishes between operating in suboptimal and supraoptimal temperatures (below or above *T_o_* respectively). The baseline moisture is used to represent the dormancy level of the seed. If *Ψ_b_* is high, the seed accumulates htu slowly for a given set of environment conditions, whereas if *Ψ_b_* is low, development is faster in the same conditions. From an initial dormancy level, *ψ_i_,* seeds lose dormancy *(Ψ_b_* becomes smaller) over time at a rate *r* that is also a function of the environmental conditions, moisture and temperature, and represents the observed process of after-ripening. *ψ_i_* is also used to represent the genetic effect on dormancy, where high *ψ_i_* represents stronger dormancy.

In Chromar, the Seed type captures information about the seed development process: Seed(gntp: (real, real), dev: real, r: real). The gntp attribute stores the genotype of the organism, *ψ_i_* (seed dormancy level) and *f_i_* (floral repression level), which is passed on to the agents representing the later stages of development and transmitted unchanged to the next generation, dev stores the cumulative development indicator (sum of htu up to the current timepoint), and r stores the after-ripening up to the current timepoint. The development rule is the following:

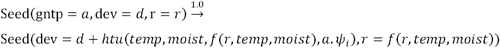

where *temp* and *moist* are fluents describing temperature and moisture. We use the ‘dot’ (.) operator for accessing the two genetic parameters of the gntp attribute. The following rule represents germination, starting the vegetative stage:

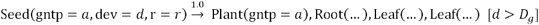

The abstract Plant agent represents the plant at the vegetative stage, along with agents for the root and the two cotyledon leaves. The initial configuration of the organs at germination is as introduced by (Chew *et al* 2014*b*). Note that the genotype attribute is passed from seed to emerged plant unchanged.

### Vegetative growth model (FM-lite) (B)

For the vegetative stage we introduce a simplified version of FMv1 (Chew *et al.*, 2014*b*) for use in studies that do not focus on circadian timing. FM-lite has three constituent models represented in Chromar with modifications to environmental responses (see below), and without the fourth, circadian clock model of FMv1.

### Timing

The timing component is the simpler flowering phenology model of (Wilczek *et al.*, 2009) rather than the augmented version in FMv1 (combination of (Chew *et al* 2012) with (Salazar *et al.* (2009) models). Vegetative development extends from dev = 0 to a threshold value, *D_f_,* where the plant flowers. The main contributing environmental factors are photoperiod, ambient temperature and vernalisation, giving the modified photothermal units, mptu, at a time *t* as:

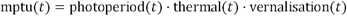

The vernalisation term accounts for both the observed requirement for a specific duration of exposure to cold and is also used to represent the genetic effect on the progress towards flowering, modelled as vernalisation(*t*) = *f(wc, f_i_*) where *wc* is the exposure to cold accumulated up to t and /_*i*_ is the genetic parameter for the initial floral repression, as in Wilczek *et al.* (2009).

In Chromar, the plant type: Plant(gntp: (real, real), dev: real, wc: real) includes the genotype attributes as noted above, the development so far (dev), and finally the accumulated winter chilling (wc). The development rule is then:

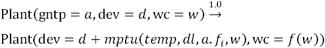

where *temp* and *dl* are fluents for temperature and day length respectively, and *w* is the present value of wc. The transition to a flowering plant, FPlant, follows:

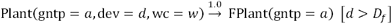

### Growth

As in FMvl (Chew *et al.*, 2014*b*), the growth component includes a carbon budget for the plant from Rasse and Tocquin (2006), which in turn includes photosynthesis rate equations based on the Farquhar *et al.* (1980) model. Growth at the organ level (rosette leaves and root) is represented based on the Greenlab model (Christophe *et al.*, 2008). We will consider a sucrose carbon pool (*c*), a starch carbon pool (*s*), and one pool for the biomass of the root and each of the rosette leaves (left panel, Figure 2). In Chromar we have the following agents to store the state (amount of carbon, or total biomass) of these pools:

- Cell(*c, s*: real) An agent that stores the amount of carbon in the sucrose (*c* attribute) and starch pools (*s* attribute). The amounts are carbon totals at the whole plant level.
- Leaf(*m*: real, *i:* int) An agent that represents a rosette leaf. It has attributes for its mass (*m*) and its index of appearance (*i*).
- Root(*m*: real) An agent that represents the root with an attribute for its mass (*m*).

**Figure 2.**
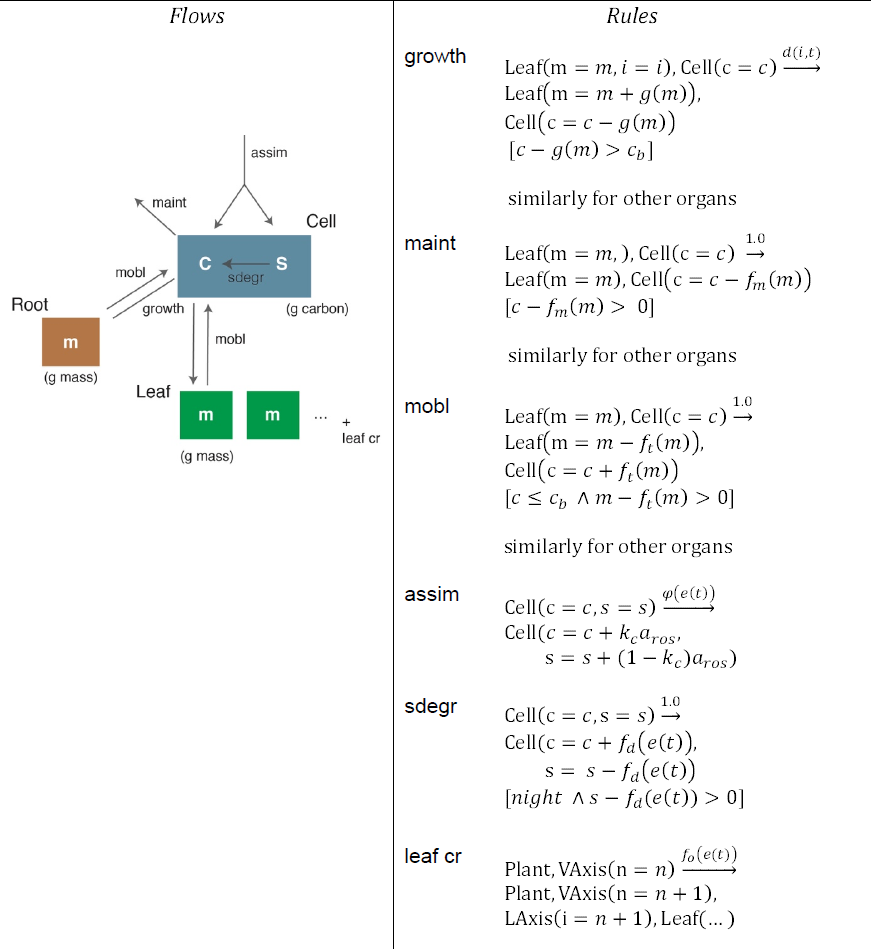
An overview of the dynamics on FM-lite (growth component of vegetative stage). The dynamics take the form of flows between different reservoirs of carbon, here shown in a graphical way (left) along with the corresponding Chromar rules (right).

For each organ we have a growth flow from the sucrose carbon pool to the mass of the organ (growth rule, Figure 2. The growth amount depends on the demand function of the organ *(d(i, t)* rule rate function) and its ‘sink strength’ *[g(m)),* which varies among organs. The value of the demand function varies over time between 1 (maximum demand) and 0 (no demand) at the end of the expansion period of the organ. The amount of carbon requested by an organ at every time unit is *g(m) · d(i, t).* Depending on the metabolic status of the whole plant (level of *c* pool) and the requests from other organs, an organ will receive either the full expected amount or a portion of it.

A flow in the opposite direction (mobl rule, Figure 2) represents carbon mobilization from the organs if the central sucrose pool (Cell(*c*)) is reduced to a critical level. Thus each organ can be either a net sink or source of carbon. For each organ, we also have a flow leaving the system from the *c* pool for the cost of the maintenance respiration and other processes of the organ (maint rule, Figure 2). Photosynthetic carbon fixation is represented by the assimilation process (assim rule, Figure 2). The amount of assimilate at every time unit is the product of the photosynthesis rate, which is a function of environmental conditions at that time step, and the projected area of the rosette. Here we use an observable, a_ros_, for the effective rosette area, which is a function of the global state of the rosette at the current time (derived from the masses of all the current leaves) and takes into account the effect of shading, as in Chew *et al.* (2014*b*). The carbon partitioning function includes a baseline partitioning to starch, then support of a target sucrose level, with excess sucrose supporting growth and a final overflow to additional starch production, as in Chew *et al.(* 2014*b*). At night, no photosynthesis occurs and carbon from the starch pools flows to the sucrose pool (sdegr rule, Figure 2). Finally, we have the creation of new leaves, which impacts the above processes indirectly by creating more demand for growth and adding maintenance costs (leaf cr rule, Figure 2). Leaves are created by the main apical meristem (VAxis agent) along with an LAxis agent that can give rise to lateral branches after flowering (see next section).

It is interesting to note that unlike FMv1 carbon partitioning between processes and organs is done explicitly whereas in our Chromar representation, partitioning is an emergent, stochastic effect of competition for the finite amount of sucrose carbon in the main reservoir. For example, partitioning of carbon among organs for growth is done explicitly in FMv1 by dividing the demand of each organ by the sum of the demands of all other organs. 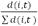 In the Chromar representation we do not have this explicit division by the global demand, which means that the amount of carbon that an organ gets is higher at each growth event but growth events are rarer because not all growth request are successful (competition). The competition therefore recovers the explicit partitioning.

#### Modifications for natural conditions

FMv1 was developed for lab conditions. As an initial approach to reflect plant responses to the broader range of relevant conditions in nature, we made the following changes:

- The rate of photosynthesis is set to 0 below 0 °C
- The maintenance cost for an organ is also 0 below 0 °C
- The rate of photosynthesis is affected by soil moisture through stomatal closure. The photosynthesis rate is affected by a stomata term *f_stom_*(*moist*) which is a simple phenomenological function that relates soil moisture and stomatal closure (France *et al.*, 1984).

These conservative changes give a lower bound on the effects of natural weather conditions.

### Comparison of FM-lite with FMvl

In addition to the weather responses, Wilczek flowering model and emergent carbon partitioning among organs, our model representation uses the stochastic rule-based Chromar as opposed to the deterministic Matlab program of FMv1. In order to compare the model representations, we simulated growth in the two models for a fixed number of hours in lab conditions, where the modifications to weather responses have no effect. The two models were simulated in lab conditions (22 °C, 12/12 light/dark cycles) for 800 growth hours and showed comparable results (Figure 3). FMv1 was simulated in Matlab while FM-lite was simulated in the Haskell implementation of Chromar and the results were averaged over five runs. The rosette mass results are the closest since they represent the development of multiple Leaf agents, masking the stochastic effects on each Leaf. The difference between the final rosette mass of the FMv1 and FM-lite (averaged over 5 runs) simulations is within 10% of the final rosette mass in FMv1. The stochasticity is more apparent for the root where the growth curves are further apart. The difference between final root mass in FMv1 and FM-lite (averaged over 5 runs) is ˜20% of the final root mass in FMv1. Sucrose carbon levels are also more variable in FM-lite, since the growth rule (removing sucrose carbon from the central pool) provides organs with a larger amount but less frequently than the small fixed amount at every time step in FMv1 (see previous section).

**Figure 3.**
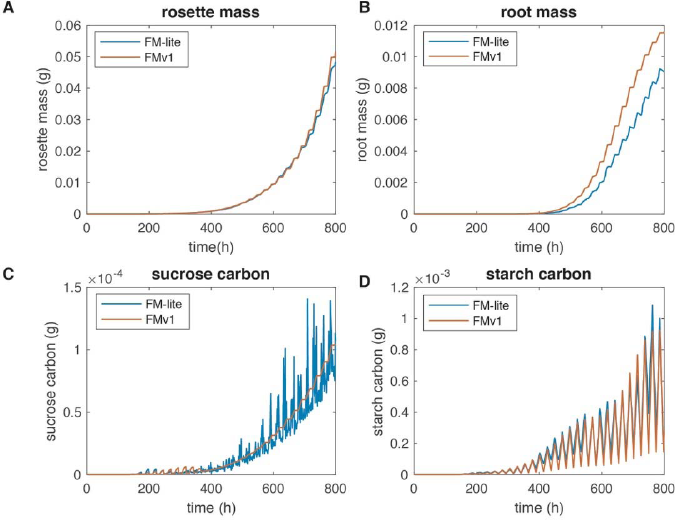
A comparison of the original FM implementation (FMvl) with the adaptation used in this work (FM-lite) for 800 hours of growth. FM-lite simulations were performed in the Haskell implementation of Chromar and results were averaged over five runs. FMvl simulations were carried out in the Matlab environment. A Comparison of simulated rosette mass trajectories between FMvl and FM-lite B Comparison of simulated root mass trajectories between FMvl and FM-lite C Comparison of simulated sucrose carbon between FMvl and FM-lite and D Comparison of simulated starch carbon between FMvl and FM-lite.

## Reproductive stage model (C)

### Timing

The timing component is a thermal time model from Burghardt *et al* (2015), representing the development of the inflorescence and seed from dev = 0 at flowering, to a threshold value, *D_s_,* where the plant disperses its seeds. Here there is no genetic input and the thermal units that accumulate at time *t* are simply the value of the temperature at *t* above a base temperature *T_b_:*

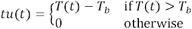

Writing into Chromar we have an FPlant(dev: real) type for a flowered plant and the following rule for its development that follows from the *tu* definition above:

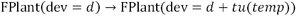

Finally, the transition to seed happens when the accumulated development reaches *D_s_:*

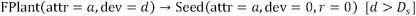

Note that the genotype attribute of the parent plant is transferred to the seeds unchanged.

### Growth

The growth component of the reproductive stage model is loosely related to the Greenlab model (Christophe *et al.*, 2008). The metabolic processes affecting the carbon budget of the plant are the same as in vegetative growth but with additional organ types to represent the Arabidopsis inflorescences. Organs appear in units (metamers) with a metamer identifier. Each growth unit on the main axis consists of an internode (stem between leaves), a leaf, and a lateral meristem that can give rise to a lateral axis. We consider only the primary axis and secondary, lateral branches, thus metamers on the lateral axis lack a further lateral meristem. All fruits on an axis are represented on its last metamer, replacing the leaf; this metamer also lacks a meristem. Two indices represent metamer position: the index of the metamer along its axis and the index of the parent metamer along the primary axis (left panel, Figure 4). We define the following new agent types to represent this structure:

- INode(*i*, *pi:* int, *m:* real) to represent the internode (stem between successive leaves). Attribute *i* is the temporal index of appearance in its axis (primary or lateral) and attribute *pi* is the parent primary metamer. The cotyledons have indices 1 and 2 on the primary axis, for example.
- LLeaf(*i*, *pi:* int, *m:* real) to represent a leaf on the lateral axes.
- Fruit(*i*, *pi:* int, *m:* real) to represent a fruit on the axis.

**Figure 4.**
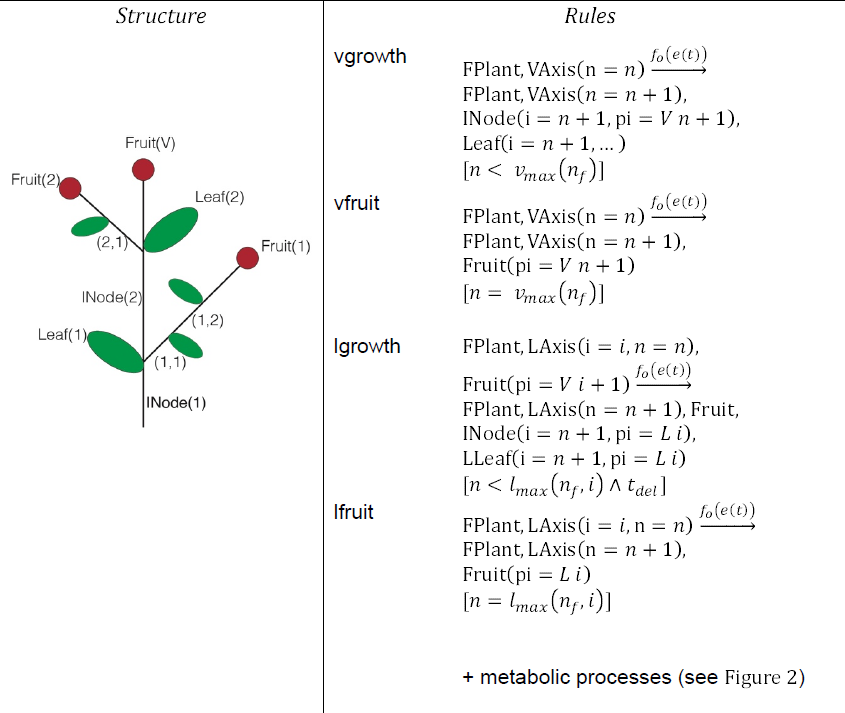
Overview of the structural part of the growth component of the reproductive stage model. The numbering scheme used to keep track of the positions of the organs in the inflorescence architecture is shown on the left. On the right the Chromar rules used to grow a structure like the one on the left panel (see main text for details).

The maximum number of inflorescence metamers on the main axis is taken to be 20% of the number of rosette leaves at flowering time (*n*_*f*_) and given by *v*_max_(*n_f_*) (Pouteau and Albertini, 2009). The maximum number of growth units on each lateral axis is given by *l*_max_(*i*), a decreasing function of the index of the lateral axis starting from a maximum of 6 at the axis after the cotyledons (index 3) and going to a minimum of 1 at the topmost lateral branch (Mundermann *et al.*, 2005). The topmost lateral axis can only appear with a delay after the apical fruit has appeared on the primary axis. Each successive lateral branch going down can only start developing with a delay after the fruit of the axis above it has appeared. The delay associated with lateral axis growth, given in the rules by *t_del_* is a function of the metabolic state of the plant, as described (Christophe *et al.*, 2008).

The new organ types have associated sink strengths and demand functions. The cauline leaves on the main axis contribute to the photosynthetically active area and can shade the rosette leaves underneath them. The lateral leaves contribute to photosynthesis without shading. Internodes and fruits do not contribute to photosynthesis. Seeds are not directly represented, so a birth function *b(m)* is required to calculate the number of seeds for a given fruit mass *m* at seed dispersal time, as described below.

### Whole life cycle model, FM-life

The Chromar framework allows us simply to concatenate the rules of timing and growth components of the three models above, to represent the whole life cycle. Then given an initial state with the genetic attributes of the plant (gntp attribute of agents) and the environmental conditions for a particular location, *e(t),* we can simulate an entire life cycle from seed to seed. The timing components of the model give us the timing within the year of the growth period (vegetative + reproductive stages) and therefore the environmental conditions that the plant is exposed to during growth. The growth components predict growth at the individual organ level with these environmental conditions and therefore give us the environmentally determined seed number given by the *b(m)* function.

### Population level model and plotting conventions

Since FM-life estimates the number of seeds at the end of the life cycle, these can initiate multiple independent copies of the model in the next generation. We then have a classical evolutionary birth process, sometimes called a branching process since it unfolds in treelike way. The potential number of individuals in generation *i, n_i_,* is equal to the sum of the number of seeds produced by the individuals in the previous generation (see Discussion). Dormant seed never die in the model and may germinate after several years (Burghardt et al.,2015).

Since we are using an individual-based model, *n*_*i*_ becomes computationally prohibitive to simulate over decades of population growth. In order to overcome this limitation, we simulated the timing (phenology) and growth components sequentially and used conservative birth functions *b(m)*. Figure 5 introduces the plotting conventions for these results. The timing components were first simulated with *b(m)* = 1, such that each plant makes one seed, as in Burghardt et al. (2015). The phenological simulation results in an unbranched sequence of developmental stage timings for each lineage (Figure 5C). The simulation results for several decades typically revealed a small number of life cycle growth strategies, from clusters of individual life cycles. The clusters were generated using *k*-means clustering, where *k* is chosen by visual inspection of the life cycle plots (Figure 5A). Alternative clustering approaches might be an area for future work. Figure 5A shows the distribution over a year of all individual life cycles that conformed to two contrasting life cycle strategies under environmental conditions for Valencia (see Results). Cluster membership depends on the dates and durations of multiple developmental stages. This is hard to visualize, because the timing of any single developmental stage partially overlaps among different strategies. Figure 5B therefore summarises the median dates of all three developmental transitions in each strategy, here illustrated by 1. a summer growth strategy and 2. a winter growth strategy. In the next stage, the growth models were simulated once per cluster, with the environmental conditions associated with the typical timing of that cluster (median vegetative and reproductive stages). This returns the typical biomass of organs over time, including the fruit mass at seed dispersal (ml for cluster 1, m2 for cluster 2; Figure 5D). Finally, each life cycle is assigned the fruit mass *m* associated with its cluster, and thereby a growth-based, reproductive success *b(m)* that evaluates to 0 in some cases. Thus, the second stage recovers a version of the branching lineage tree, where some lineages die out (Figure 5E).

**Figure 5.**
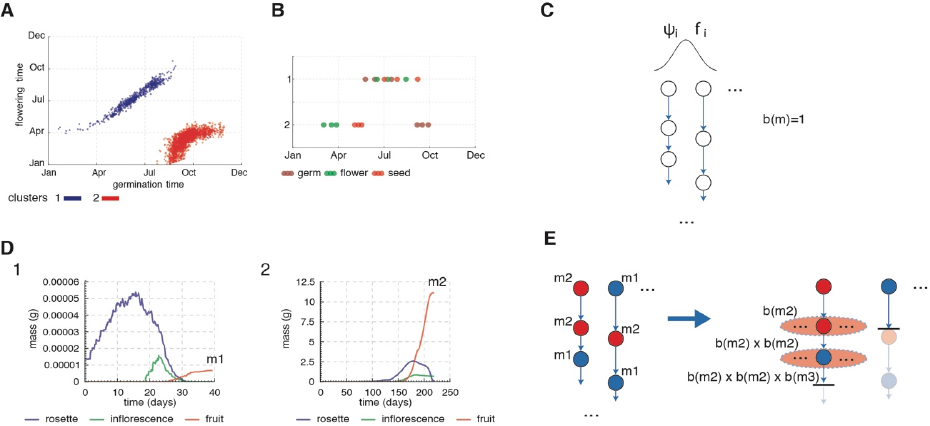
The two-stage simulation of the population level model. A The distributions of developmental events (germination, flowering) from the phenology-only simulation of the population model (C) with the identification of two clusters representing two distinct strategies. B The 25-th, 50-th, and 75-th percentiles of the distributions of developmental events of the two clusters from A. C Illustration of the phenology only simulation with b(m)=1 (each plant makes a single seed) D Results of simulations of the growth models for the median dates of developmental events (B) E Illustration of the assignment of fruit masses to the lifecycles of the phenology-only simulation (C) according to their clusters. This recovers the full branching population process.

Our output population measure is the total population of plants over all lineages over all generations. For example, consider a lineage with three generations starting with a plant with final fruit mass *m*_11_. For the next generation we have *b*(*m*_11_) and then *b*(*m*_11_) × *b*(*m*_21_). The population measure for that lineage is 1 + *b*(*m*_11_) + *b*(*m*_11_) × *b*(*m*_21_). The population measure for multiple lineages starting from multiple plants in the initial population is the sum of the population measures of all the lineages. This requires a birth function, which we use in a very simple form, as follows:

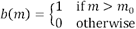

A plant produces one seed or none, the latter in life cycles with fruit mass at seed dispersal *m* less than a threshold *m_0_.* Below, we make some conservative choices for the value of the reproductive threshold value, *m*_0_, to explore the effect on the output population measure.

Finally, we distinguish three sources of variability in the population model: (i) weather varies between years, (ii) genetic parameters can vary among the initial population if their values are chosen probabilistically, and (iii) simulation results vary due to stochasticity in the model representation.

### Weather data

For the phenology model simulation we used the weather data that accompanied the Burghardt et al. (2015) model, available from a Dryad repository (Burghardt et al. 2014). In this dataset weather inputs over 60 years were generated stochastically for four locations in Europe: Halle, Valencia, Norwich, and Oulu. The weather inputs include values for temperature, moisture, and daylength.

For the growth simulations we used weather data from the ECMWF ERA-Interim dataset over the years 2010-2011 (Dee *et al.*, 2011). A program was used to generate hourly inputs given daily averages from the dataset for temperature and radiation. For the soil moisture input used in the photosynthesis rate calculation we used a daily average of soil moisture values from the dataset and assumed that is constant throughout the day (swvl parameters in the ERA dataset).

## Results

The population of FM-life models (see Methods) allows us to test how growth processes that alter reproductive success affect the life history strategies of Arabidopsis growing in different environmental conditions (location) and with different genetic parameters in the initial population. We can therefore explore the genotype x environment interaction, using a population measure. To illustrate this potential, we compare simulation results for two previously-studied locations, Valencia (Spain) and Oulu (Finland), and two opposing combinations of genetic parameters, high seed dormancy /high floral repression (HH) and low seed dormancy /low floral repression (LL). Within an initial population of 100 seeds, the seed dormancy levels, *ψ_i_* were assigned probabilistically, sampling from a normal distribution with mean 0.0 and standard deviation, 1, for the Low dormancy level (L) and mean 2.5 with the same standard deviation for the High dormancy case (H). Floral repression was fixed at either 0.598 for the Low level (L) and 0.737 for the High level (H), values that were chosen to reflect the behaviour of natural populations of Arabidopsis in Wilczek *et al* (2009). Both parameter choices follow Burghardt et al. (2015). The simulation time period was 60 years and, as in Burghardt *et al* (2015), we discarded the first 15 years of the simulation to focus on stable life history strategies. A key difference from the earlier work is that even our conservative choice of birth function (see Methods) allows some lineages to die out.

## Valencia

Figure 6 shows the results of the two-stage simulation for a population of the LL genotype in Valencia (Figure 6A). We identify four possible life history strategies based on the timing (phenology) components of the FM-life model:

1. *summer-only strategy* where the entire growth is in the summer. The growth period is quite short and the conditions unfavourably hot and dry. In the growth simulation, the rosette leaves senesce before the reproductive stage (blue curve). The drought effect on photosynthesis severely limits the carbon available for fruit mass (red curve).
2. *spring strategy* where the entire growth period is in the spring. The growth period is only slightly longer than the *summer-only* strategy but it falls in more favourable weather conditions. The rosette lifetime extends beyond flowering to support fruit growth, which combined with favourable weather gives high fruit mass.
3. *winter-repr strategy* spans the winter/early spring period. A short vegetative period in the end of summer/early Autumn ends with flowering and a long reproductive stage over the winter/early spring. The rosette is senescing when favourable conditions return in early spring, seriously limiting fruit development.
4. *winter-veg strategy* again spans the winter/early spring period. The life cycle duration is similar to strategy 3 but slightly later germination delays flowering until Spring. The rosette grows all winter, overlapping with a short reproductive stage and supporting high fruit mass.

**Figure 6.**
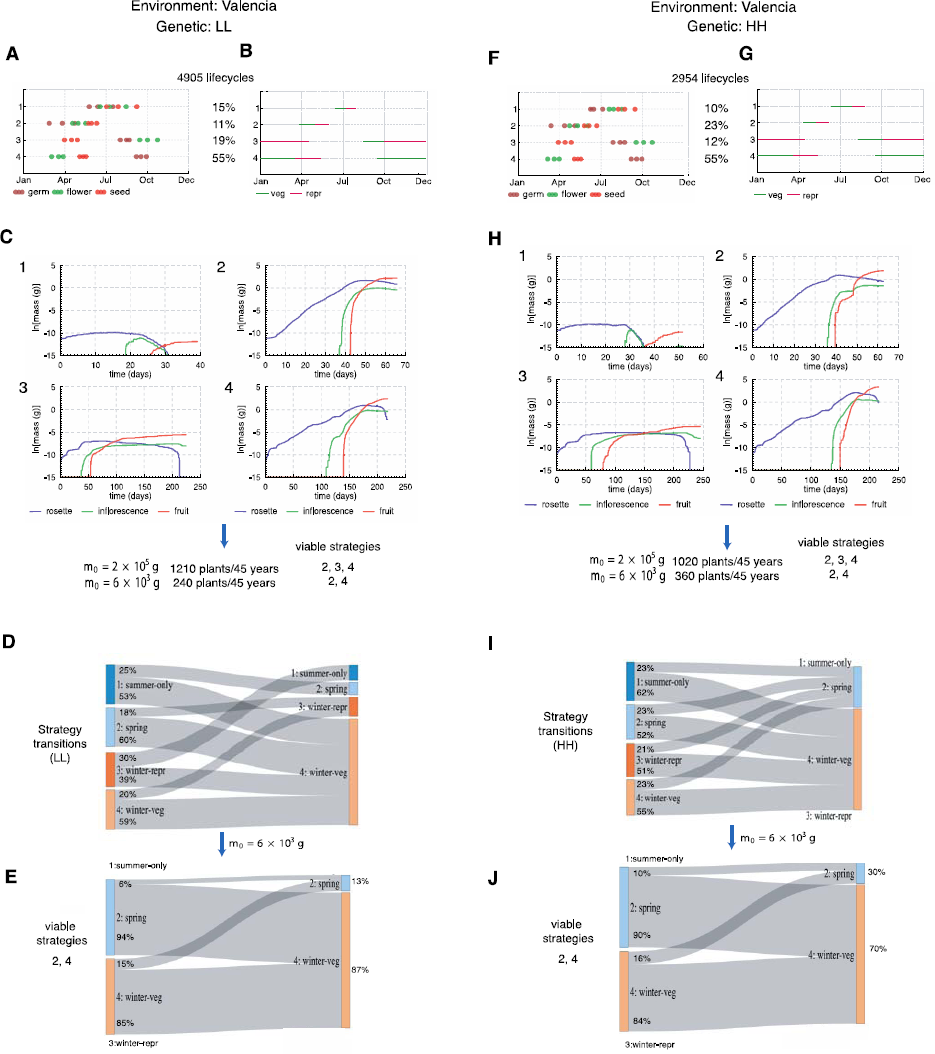
Population experiments in Valencia in two different genetic backgrounds (LL − low dormancy, Low floral repression and HH-High dormancy, High floral repression). A Results of the phenology-only simulation results for the LL genotype. The 25-th, 50-th, and 75-th percentile of the distribution of developmental events are shown for each identified cluster B Illustration of growth stages over a year for each cluster from A according to the median time of the distribution of developmental events for each cluster. C Growth simulations over the growth period shown in panel Bfor each cluster. D Probabilities of successive strategies. E Probabilities of successive strategies after eliminating life cycles with strategies 1 and 3 using a reproductive threshold. F, G, H, I, J Equivalently for the HH genotype in the same location.

Plants with life cycle strategies 2 and 4 predicted orders of magnitude more fruit mass than plants with life cycle strategy 3 and or the least successful strategy 1 (Figure 6C). This result clearly ranked the strategies available to plants of the LL genotype, although the absolute values of the predicted biomass are less certain (see Discussion). The 100 plants amassed 4905 potential lifecycles over 45 years of phenological simulation (Figure 6A).

Without a minimum mass threshold *[mo]* for reproduction, 66% of potential life cycles followed the more successful *spring* and *winter-veg* strategies (2 & 4; Figure 6C). Figure 6D shows the sequential transitions between strategies. For example, 60% of potential plants following the successful *spring* strategy (2) disperse their seeds early enough for the next generation to adopt the *winter-veg* strategy (4), achieving two generations per year. These transitions underlie the bimodal distribution of life cycle times reported by Burghardt et al. (2015) for this simulation.

Simulation of the HH genotype (Figure 6F, 6J) identified similar strategies. Since the seed have longer dormancy, the population amassed fewer potential life cycles (2954 as opposed to 4905 in the LL case; Figure 6F). The growth and final fruit masses are different because of slight variation in timing of the growth period but strategies 2 and 4 are again more successful than strategies 1 and 3 (Figure 6H). A higher fraction of potential life cycles followed the successful strategies (78% as opposed to 66% in the LL case; Figure 6G). Higher seed dormancy reduced the germination in the summer and early autumn that led to the less successful strategies 1 and 3, so any strategy was likely to be followed by either strategy 2 or 4 in the next generation (Figure 6).

In order to calculate the population success we make two choices for the reproduction mass threshold, *m_0_,* which eliminate one or both of the least successful strategies. Choosing a value *m_0_* = 2 x 10^−5^ g (the mass of a single seed) eliminated the *summer-only* strategy from both genotypes, which gives a population of 1210 plants over 45 years in the LL case (Figure 6C). The HH genotype allows a larger percentage of viable life cycles but we predict fewer plants (1020) since the number of potential life cycles was lower (Figure 6H). Choosing a value *m_0_ =* 6 x 10^−3^ g left only two viable strategies, 2 and 4, for both genotypes. Reciprocal transitions between the strategies were still possible but *winter-veg* was strongly favoured (Figure 6E, 6J). The LL genotype predicted 240 plants in total over 45 years, compared to 360 plants for the HH genotype: G x E interaction favoured the HH genotype despite its smaller number of potential life cycles. Thus, modelling the growth processes not only distinguished among the potential life cycle strategies within a genotype but also between the genotypes.

### Oulu

The equivalent simulations were performed for conditions in Oulu, Finland in the same LL and HH genetic backgrounds (Figure 7). The results indicated 3 potential life cycle strategies (Figure 7A, 7B, 7E, 7F):

1. *summer-only strategy* where the entire life cycle occurs in the summer. The vegetative period is short, the rosette is very small and supports negligible fruit growth (Figure 7C).
2. *winter-repr strategy* where a life cycle of almost a year has a very short vegetative stage, followed by a long reproductive stage over the winter. Again, the very small rosette supports little fruit growth in the following Spring.
3. *winter-veg strategy* where the plant over-winters in the vegetative stage. Unlike in Valencia, the rosette grows little over the winter. Rapid rosette growth in the following spring supports a substantial inflorescence and fruit development, though the predicted fruit mass is smaller than in Valencia.

**Figure 7.**
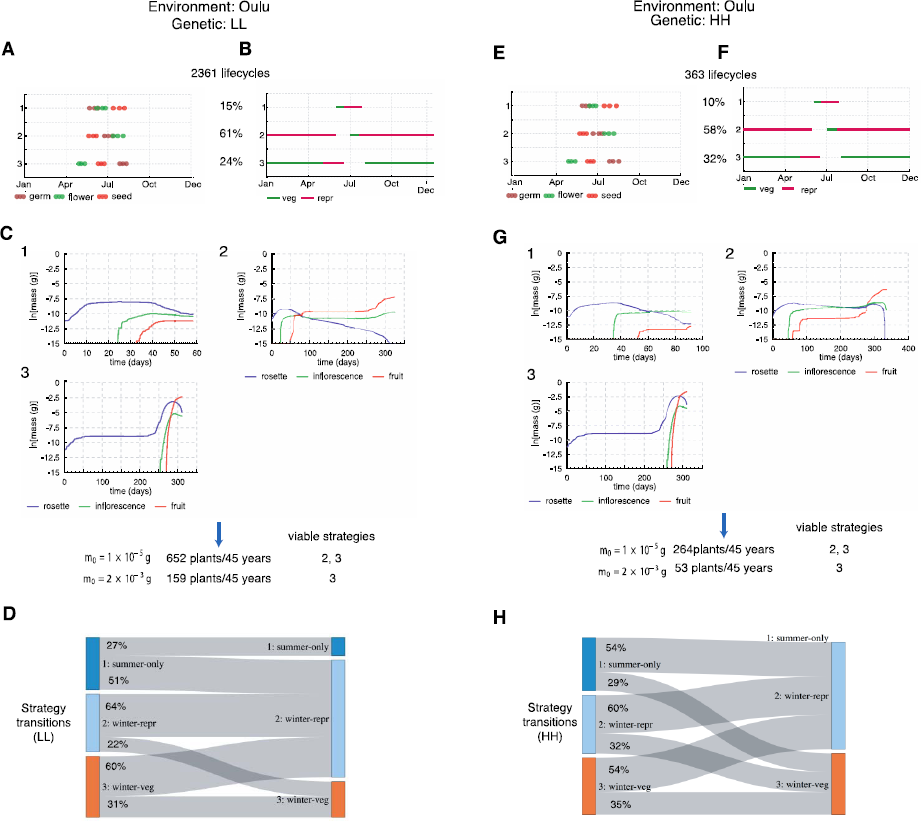
Population experiments in Oulu in two different genetic backgrounds (LL − low dormancy, Low floral repression and HH-High dormancy, High floral repression). A Results of the phenology-only simulation results for the LL genotype. The 25-th, 50-th, and 75-th percentile of the distribution of developmental events are shown for each identified cluster B Illustration of growth stages over a year for each cluster according to the median time of the distribution of developmental events for each cluster. C Growth simulations over the growth period shown in panel Bfor each cluster. D Probabilities of successive strategies. E, F, G, HJ Equivalently for the HH genotype in the same location.

The severe winter conditions limited the number of potential life cycles to 2361 for the LL genotype or 363 for HH. A higher proportion of HH life cycles followed the successful *winter-veg* strategy (3; 32% against 24% in LL; Figure 7B, 7G). Surprisingly, a majority of life cycles for both genotypes followed the *winter-repr* strategy (2). Applying the reproductive threshold mass, *m_0_,* eliminated one or both of strategies 1 and 2 (Figures 7C, 7G), suggesting a strong selective pressure for greater floral repression to reduce the number of *winter-repr* life cycles. With *m_0_ =* 2 × 10^−3^ g, the LL genotype yielded 159 plants over 45 years compared to 53 plants for HH. All G x E combinations had actively-growing plants at the end of the simulation. Interestingly, plants of the HH genotype had higher average reproductive success per plant in Oulu yet the LL plants were more successful by our population measure. The faster development of LL plants allowed more, short lifecycles within the simulated interval (consistent with the phenology model alone).

## Discussion

We present a whole-life-cycle multi-model for growth and reproduction of *Arabidopsis thaliana,* FM-life, combining phenology models that time the developmental stages and growth models to predict organ biomass. The simple, FM-lite model of vegetative growth, and its extension to the reproductive stage in FM-life, simulate broader, mechanistically-founded components of fitness at the individual plant level compared to the phenology models alone. Most insights from the component models naturally remain (Rasse and Tocquin, 2006; Christophe *et al.*, 2008; Wilczek *et al.*, 2009; Burghardt *et al.*, 2015). Multi-models are helpful in emphasising interactions. The cauline leaves in our inflorescence model, for example, extend the duration of photosynthetic competence. As cauline leaves can be produced 6 months later than early rosette leaves in the *winter-veg* strategy (Figure 7), they remain active photosynthetic sources (Earley *et al.*, 2009; Leonardos *et al.*, 2014) when the rosette leaves are senescing. The growth models provided the fruit mass that we used as an indicator of reproductive success, such that metabolic and developmental processes of growth informed a more mechanistic understanding of ecological, population dynamics over multiple generations.

The growth model allowed us to discriminate among alternative life cycle strategies in each G × E combination, by selecting against strategies that were compatible with the phenology models alone but had qualitatively worse growth. In previous work, strategies with high seed dormancy in southern Valencia and low dormancy in northern Oulu were noted to align with the behaviour of the cognate wild populations (Atwell *et al.*, 2010; Chiang *et al.*, 2011; Méndez-Vigo *et al.*, 2011; Burghardt *et al.*, 2015). In each GxE combination, individual plants in our simulations might adopt alternative life cycle strategies. The less-successful strategies were lethal in our model, eliminating >95% of potential lifecycles (simulated by the phenology model alone) for the LL genotype in Valencia, for example (Figure 6A, 6C). Thus, our results supported the observed genotypic distinction between Valencia and Oulu, because the requirement for a minimum fruit mass eliminated more lineages of the less-successful genotype in each case (Figures 6C, 6H and 7C, 7G).

Our approach might appear conservative, as the binary birth function (one seed/no seed) ignored variation in seed mass among life cycle strategies, which might otherwise reinforce the advantage of successful strategies. The successful genotype LL in Oulu, however, had lower reproductive success per plant than HH, suggesting a more subtle balance of advantage. Genotypes with Low dormancy and High floral repression (LH) are observed in far northern locations (Atwell *et al.*, 2010). We therefore simulated the LH genotype (Figure 8). LH plants delayed flowering time enough to reduce the frequency of potential *summer-only* life cycles to 9% compared to 15% in LL (Figure 8B) and increased the fruit mass of the successful *winter-veg* life cycle close to the HH genotype (Figure 8C). The LH model predicted slightly higher reproductive success overall, returning 171 life cycles (Figure 8C) compared to 159 for the LL variant (Figure 7C), consistent with the observation of LH genotypes at this location.

**Figure 8.**
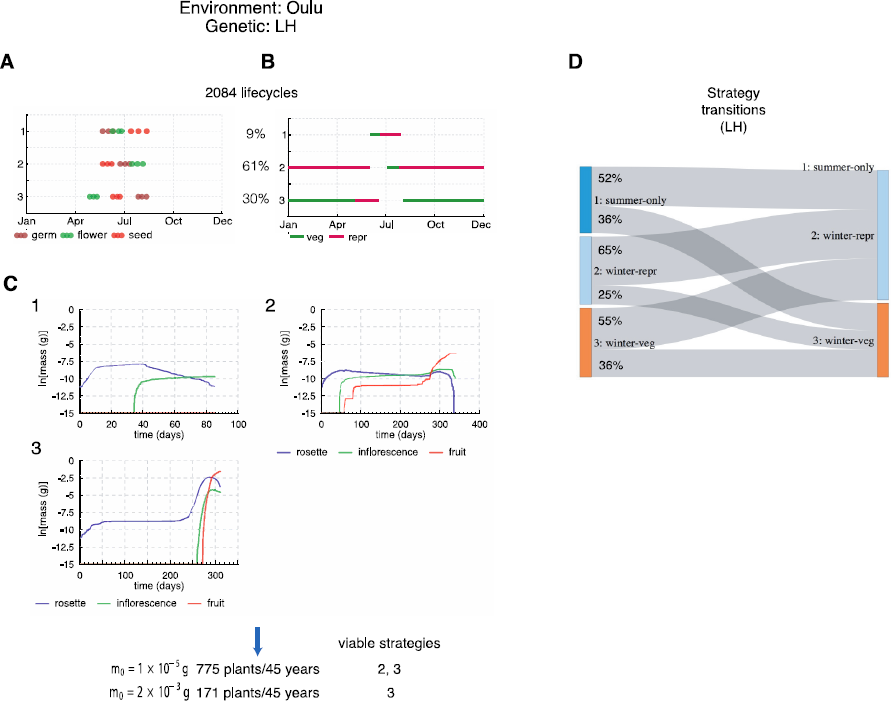
Simulation results for a combined variant in Oulu (LH − low dormancy, high floral repression). A Results of the phenology-only simulation results. The 25-th, 50-th, and 75-th percentile of the distribution of developmental events are shown for each identified cluster B Illustration of growth stages over a year for each cluster according to the median time of the distribution of developmental events for each cluster. C Growth simulations over the growth period shown in panel Bfor each cluster. D Probabilities of successive strategies.

A limitation of our work arises from the fact that the phenology component models of FM-life have been validated against field data (Wilczek *et al.*, 2009; Burghardt *et al.*, 2015) whereas the growth component models have not (Rasse and Tocquin, 2006; Christophe *et al.*, 2008). Biomass simulations are inevitably sensitive to the timing of the growth period, because a longer interval of exponential growth in good conditions rapidly changes absolute biomass, as illustrated in Figure 9D. Nonetheless, the FM-life model predicted unreasonably high fruit mass in some cases. The binary birth function ensured that this had no effect on our population measure. Among possible gaps in understanding of the environmental effects on growth in natural settings or in our representation, we repeat our previous caution (Chew *et al.*, 2014*b*, 2017) that models of nutrient balance for Arabidopsis will be helpful. Rosette biomass in the Framework Model is understandably sensitive to photosynthetic parameters (Chew *et al.*, 2014*b*) yet these have not been validated in Arabidopsis across the wide range of photoperiods and temperatures simulated here (Walker *et al.*, 2013). FM-life predicts a discretised fruit mass and hence reproductive success for a typical representative of each life cycle strategy, approximating an underlying, continuous distribution of fruit mass. The accuracy of this approximation will depend on the variation within clusters. The benefit lies in computational tractability, allowing us to simulate differential reproductive success that is informed by understanding of growth processes.

**Figure 9.**
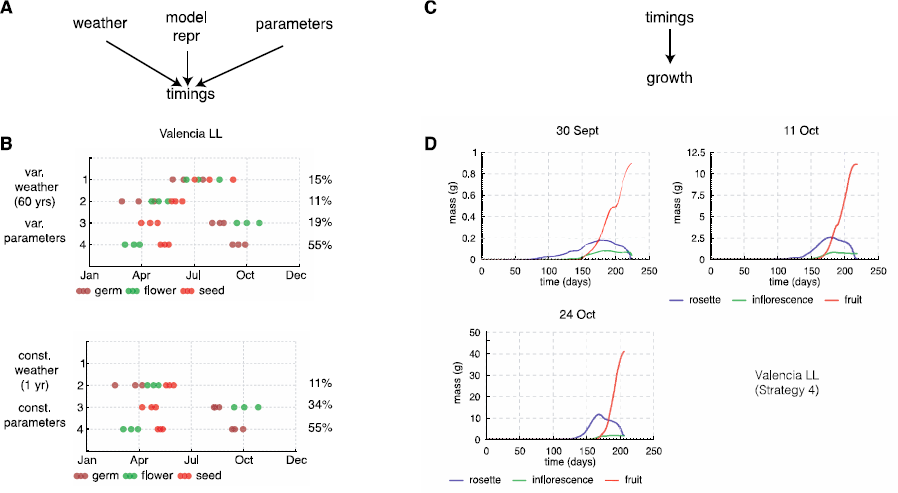
Robustness of timing and growth simulation results. A Sources of variability for the timing results. B Timing results for variable weather (60 years) and variable genetic parameters chosen from a distribution and timing results for simulations with constant weather (same 1-year weather over 60 years) and constant parameters. The timings of the strategies are very similar but strategy 1 life cycles have moved to strategy 3. This could be because of the particular year of weather data we used for the simulations. C Sources of variability for growth results D Growth simulation results starting from three different germination dates (corresponding to 25-th, 50-th, and 75-th percentiles of the distribution of germination times) of strategy 4 in Valencia LL (Figure 6)

Our approach builds upon previous models that predict fitness and population processes in Arabidopsis, which have focussed on developmental components of fitness or on phenology (Prusinkiewicz *et al.*, 2007; Satake *et al.*, 2013; Springthorpe and Penfield, 2015). Linking these components sharpens ecological insight, by understanding the performance of genetic variants in the environment that underlies differences in fitness (see discussions in (Donohue *et al.*, 2015; Doebeli *et al.*, 2017)) and can thus inform evolutionary hypotheses. Adding genetic variation between generations will in future model Arabidopsis evolution explicitly, perhaps after competing genetic variants *in silico* using adaptive dynamics approaches (Brännström *et al.*, 2013; Weiße *et al.*, 2015). Thus, the FM-life model offers a further tool to bridge among disciplines in plant biology, ecology and evolution.

**Table 1.** *Table of relevant parameter values. Developmental thresholds for the timing components of the three stages are from the Burghardt et al. model*

| Name | Units | Value | Description |
| --- | --- | --- | --- |
| $D_g$ | °C MPa | 1000 | Developmental threshold for germination |
| $D_f$ | °C h | 2604 | Developmental threshold for flowering |
| $D_s$ | °C | 8448 | Developmental threshold for seed dispersal |
| $m_0$ | $g$ | various | Reproductive threshold |
| $\psi_i$ | MPa | Low(L)=0.0<br>High(H)=2.5 | Initial seed dormancy (distribution means) |
| $f_i$ | None | Low(L)=0.598<br>High(H)=0.737 | Initial floral repression |

## Acknowledgements

We are grateful to L. Burghardt and K. Donohue (Duke University) for sharing data and model definitions during a post-publication embargo period, and to L. Smallman and M. Williams (University of Edinburgh) for expert assistance with meteorological inputs and eco-physiological simulation. Argyris Zardilis was supported by an iCASE studentship from the Biotechnology and Biological Sciences Research Council (award BB/K011294/1) with Simulistics Ltd.

